# Sonographic Analysis of Abscess Maturation in a Porcine Model

**DOI:** 10.1101/2020.09.28.317321

**Authors:** Daniel F. Leotta, Matthew Bruce, Yak-Nam Wang, John Kucewicz, Tatiana Khokhlova, Keith Chan, Wayne Monsky, Thomas J. Matula

**Affiliations:** Applied Physics Laboratory, University of Washington, Seattle, WA 98105, USA; Department of Gastroenterology, University of Washington, Seattle, WA 98105, USA; Department of Radiology, University of Washington, Seattle, WA 98105, USA

**Keywords:** Abscess, Ultrasound, Three-Dimensional, Curvature, Shear Wave Elastography, Doppler

## Abstract

Abscesses are walled-off collections of infected fluids that often develop as complications in the setting of surgery and trauma. Abscess care depends on size, location, composition and complexity, among other patient factors. The goal of this work is to describe, using the latest ultrasound imaging technologies, the progression of abscess development in a porcine animal model. Intramuscular or subcutaneous injections of bacteria plus dextran particles as an irritant led to identifiable abscesses over a 2- to 3-week period. The abscesses were imaged at least weekly with B-mode, 3D B-mode, shear-wave elastography (SWE) and color flow imaging. Mature abscesses were characterized by a well-defined core of varying echogenicity surrounded by a hypoechoic capsule that was highly vascularized on Doppler imaging. Size and shape changes during development were quantified with 3D imaging. With SWE, the lesion stiffness varied interiorly and generally decreased over time. These ultrasound features potentially provide biomarkers to facilitate improved selection of treatment strategies for abscesses.

## Introduction

Infected abscesses are walled-off collections of inflammatory cells, cellular debris, and bacteria. They often develop as complications in the setting of surgery, trauma, systemic infections, and other disease states. Abscess care depends on size, location, composition and complexity, among other patient factors. For deeper abscesses, current treatment is typically limited to antibiotics with long-term catheter drainage, or surgical washout when inaccessible to percutaneous drainage or unresponsive to initial care efforts. Ultrasound imaging can assist in reliable identification of abscesses so that appropriate treatment is applied (O’Rourke et al. 2015). For example, ultrasound can be used to distinguish abscesses from other soft-tissue pathologies such as cysts, hematomas, phlegmons and cellulitis. It can also be useful in planning therapy strategies by establishing extent, maturity, and structural features such as loculations (internal chambers).

Variations in echogenicity observed within abscesses have been noted in B-mode imaging studies, which can make definitive diagnosis challenging (Kuligowska et al. 1982). Contrast ultrasound studies have been used to distinguish between inflammatory tissue that may represent early infections (phlegmons), and fully developed mature abscesses by measuring enhancement within the infected tissue (Ripollés et al. 2011, 2013). Full enhancement within the tissue signifies a phlegmon, while lack of internal enhancement is a sign of a mature abscess containing potentially-drainable fluid. Blood flow surrounding the core of an abscess has also been reported with power Doppler imaging (Arlsan et al. 1998) and with contrast-enhanced ultrasound (Liu et al. 2008). Elastography has also been reported as a potential modality to diagnose and characterize abscesses (Gaspari et al. 2009).

Both B-mode and contrast imaging have been used to classify abscess maturity by comparing imaging features observed in abscesses across different patients (Kunze et al. 2015). However, the early progression of abscess development as observed by ultrasound has yet to be fully described. The objective of this study was to use the latest advances in non-contrast ultrasound technologies to characterize abscess development in a novel porcine animal model (Wang et al. 2020). Ultrasound imaging was performed at least weekly to evaluate abscess maturation following intramuscular injection of bacteria. The controlled induction of the abscess allowed for imaging at definitive time points in abscess development. Abscesses were imaged during the maturation process over 4 weeks, using B-mode, 3D, elastography and Doppler ultrasound, to evaluate changes in echogenicity, size, shape, stiffness and blood flow.

## Materials and Methods

### Animal Model

All procedures were approved by the Institutional Animal Care and Use Committee (IACUC) at R&R Rabbitry (Stanwood, WA). Two female domestic swine (Yorkshire/Hampshire cross, age 3-6 months, weight 45-65 kg) were used in this study. In a recent paper (Wang et al. 2020) we described the development of a porcine animal model in which multiple large multiloculated bimicrobial abscesses can be formed at distinct sites in the same animal. These abscesses satisfy the true definition of an abscess and are formed within a short period. Briefly, on the day of inoculation, the bacterial mix, consisting of E. coli (strain ATCC 25922, American Type and Culture Collection, MD, USA), B. fragilis strain (ATCC 23745, American Type and Culture Collection, MD, USA) and sterile dextran microcarrier bead suspension in a ratio of 1:1:2 was prepared so that the final inoculum contained approximately 106 cfu/ml E. coli and 10^6^ cfu/ml B. fragilis. Bacterial concentration was determined using spectrophotometric absorbance readings and confirmed using Compact Dry™ EC100 and TC assay plates (Hardy Diagnostics, Santa Maria, CA USA).

Animals were sedated (Telazol/Ketamine/Xylazine) and maintained under a surgical plane of anesthesia with isoflurane. All animals were instrumented to monitor heart rate, ECG and blood oxygen saturation and temperature. The haunches, abdomen and in one animal the neck were depilated, cleaned and sterilized with consecutive rounds of isopropyl alcohol. Injections were performed under ultrasound guidance into 6 to 8 separate sites in a single animal: 1.5 cm deep into the left and right biceps femoris, brachiocephalicus, and subcutaneously on either side of the midline. Before the intramuscular injections, the tip of the needle was moved back and forth approximately 1 cm fifty times, in a fanning motion, to create a small pocket of injury before injecting. For each animal 10 ml of the inoculate were injected at each site and the location was marked with dye. After the inoculations, the animals were recovered and monitored every day for signs of pain or distress, reduction in appetite, loss of weight and mobility. Abscesses were allowed to form over 3-4 weeks.

### Ultrasound imaging of abscesses

Growth and development of a total of 14 abscesses in the two animals were monitored at least weekly with ultrasound imaging using an Aixplorer (Supersonic Imagine, Aix-en-Provence, France) with a 4-15 MHz linear probe. Modalities used included B-mode (for general and 3D imaging), color and power Doppler (to evaluate vascularity), and shear wave elastography (SWE, to evaluate stiffness). In some cases, palpation with the ultrasound probe was used to observe movement of the abscessal contents as a gross indication of the liquidity of the pus. For one animal three-dimensional imaging was performed by tracking the ultrasound scanhead tracked with a magnetic tracking system (trakSTAR, NDI, Waterloo, ON, Canada). Custom software recorded the 3D location and orientation of a series of 2D image planes acquired while scanning the abscess, and a 3D volume was reconstructed from the 2D images (Leotta et al. 2018).

### Image analysis

Detailed image analysis was performed on two abscess cases, to first illustrate the general characteristics of a mature abscess, and then to quantify the changes in size, shape and composition during maturation. The first case is an intramuscular injection that progressed to a compact abscess over the course of 3 weeks. The second case is an intramuscular injection in a second animal that progressed to an elongated abscess over the course of 4 weeks.

The first case (ABS1) was scanned with B-mode and Doppler up to 3 weeks post-injection, and at 3 weeks it was representative of a mature abscess. Doppler imaging with plane-wave directional color power imaging mode was used to visualize the blood vessels within the abscess. The plane-wave Doppler mode (Angio PLUS) provides increased sensitivity to low velocity flows. A 3D reconstruction of the Doppler images was generated from a slow untracked sweep across the full extent of the abscess, assuming a linear scan at constant speed. A total of 605 2D images were acquired during this sweep. Since the full abscess could be visualized in a single 2D field of view for this smaller abscess, the sweep distance was determined from size measurements derived from 2D images. The color pixels on each image were automatically segmented from the background so that the 3D reconstruction represents the Doppler component only.

The second case (ABS2) was scanned in B-mode at weeks 2, 3 and 4 with the 3D image capture system. These tracked scans were used to create gray-scale volume reconstructions by compiling the series of 2D images in a regular 3D grid (Leotta and Martin 2000). The total number of images collected were 1212, 1783, and 341 for the scans at weeks 1 through 3, respectively. Manual image segmentation in a series of axial planes was performed with custom software (Leggett et al. 1997) to construct surface mesh models of the outer capsule and the inner core (Leotta et al. 2001). The surface models were analyzed with the MeshLab open-source analysis package (Cignoni et al. 2008) to quantify the dimensions (length, width, height) and volume of both the total abscess size (outer extent of the capsule) and the core. As a measure of the surface complexity, MeshLab was also used to calculate the mean curvature of the surface reconstruction of the abscess core at each time point. For each surface model. the local curvature is calculated at each mesh vertex using the method described in (Meyer et al. 2003). Curvature is zero for a flat surface, positive for a convex surface, and negative for a concave surface; increasing magnitude signifies greater deviation from a plane.

For Case ABS2, shear wave elastography was performed at weeks 1 through 4. At weeks 2 and 3, slow sweeps at constant speed were acquired along the length of the abscess, imaged in cross-section. The 3D tracking system was not used during these scans. Instead, 3D reconstructions were approximated by compiling the image series into a 3D volume assuming a linear scan at a constant sweep speed. The scan distance was determined from the 3D reconstructions of the tracked B-mode scans. The total number of images compiled for each scan was 862 and 1584 for weeks 2 and 3 respectively. SWE values for the abscess core were extracted by 1) isolating the core in the 3D volume by manual segmentation, and 2) converting the color pixels within the core to stiffness values in kPa according to the SWE color map.

Because this abscess was close to the skin, it was also possible to use probe palpation to visualize the motion of lower-viscosity liquid components under B-mode when the abscess was pushed on by the ultrasound probe. A video clip was acquired at week 3 post-injection during several cycles of manual compression of the abscess. From a portion of the video, particle tracking was used measure the motion of the liquid pus within the core. For this analysis, particle displacement direction and magnitude were tracked over 13 sequential video frames using optical flow analysis (Fleet and Weiss 2006) as the probe pushed on the abscess. Optical flow is a video processing method for tracking movement within a video sequence based on spatial and temporal image gradients. The Lucas-Kanade optical flow method was used, which uses least-squares regression to solve the optical flow equation for regions of pixels rather than individual pixels (Lucas and Kanade 1981).

Doppler images were also acquired each week for this case. Doppler modes included standard color flow, directional color power, and plane-wave directional color power. The plane-wave Doppler mode provides increased sensitivity and resolution compared to conventional Doppler imaging, providing enhanced visualization of small vessels and low-velocity flow.

### Histopathology

Histological analysis was performed for case ABS2 at the end of the 4-week study period. After the animal was euthanized, the abscess was carefully removed *en bloc* and fixed in 10% neutral buffered formalin for histological evaluation. The fixed lesion was grossed and processed to maintain the lesion as whole cross-sections as much as possible. Serial sections were stained with Hematoxylin and Eosin (H&E) for general tissue morphology, Masson’s trichrome stain to visualize the presence of a fibrous capsule, or labeled with anti-CD31 antibodies to visualize blood vessels. The lesion was evaluated for the presence of a connective tissue capsule; the extent of the capsule; the presence of infected material and cell debris; and blood vessels.

## Results

### Abscess maturity

For all 14 abscesses imaged for this study the initial bacteria inoculation led to a pocket with an identifiable capsule at about 2-3 weeks. Figure 1 shows abscess case ABS1 at 3 weeks post-injection. It shows the abscess core surrounded by a fibrous capsule. Doppler imaging shows a dense network of blood vessels within the capsule. The 3D distribution of the blood vessels in the capsule is shown in Figure 2 at 2.5 weeks. This abscess is representative of the general features that were visualized at maturation: a well-circumscribed, avascular, mixed echogenic core surrounded by a vascularized hypoechoic capsule.

**Figure 1.**
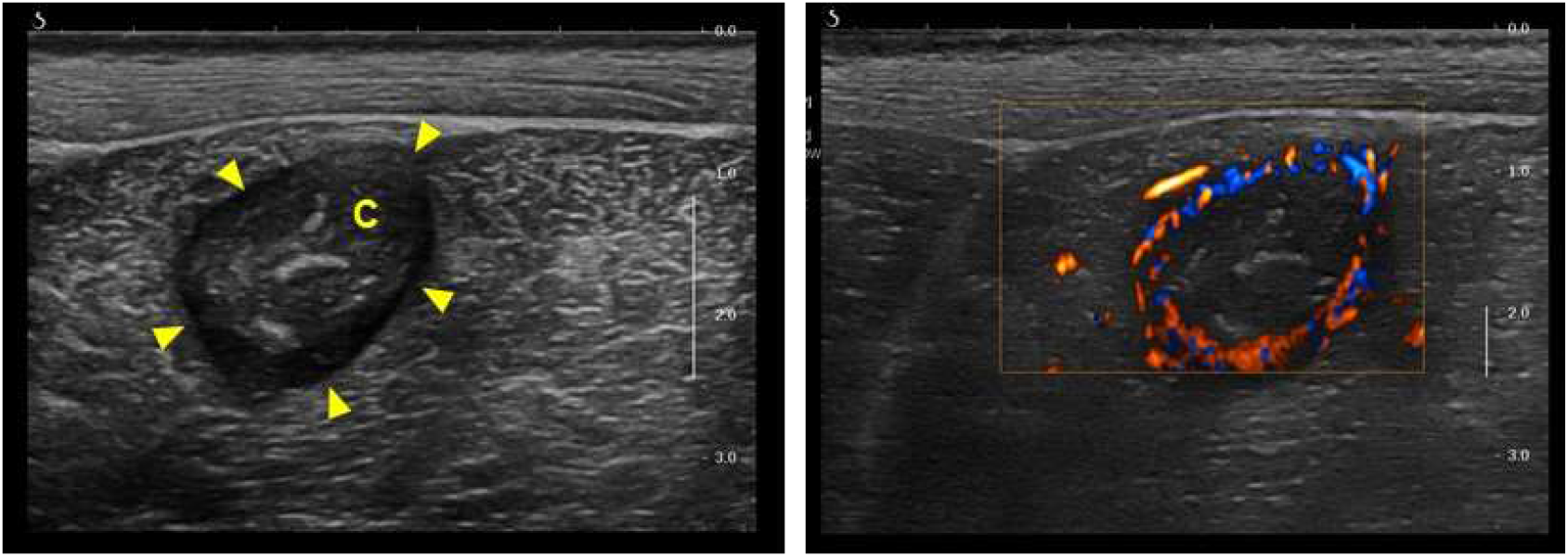
Representative B-mode (left) and color Doppler (right) images for intramuscular abscess ABS1 imaged at 3 weeks post-injection. The center of the abscess lies approximately 2 cm below the skin surface within the biceps femoris. The B-mode image demonstrates the salient features of a mature abscess, including a well-circumscribed hypoechoic capsule (arrow heads) and a core (C) with variable echogenicity. Plane-wave directional-power color Doppler shows an avascular core surrounded by a highly vascularized capsule.

**Figure 2.**
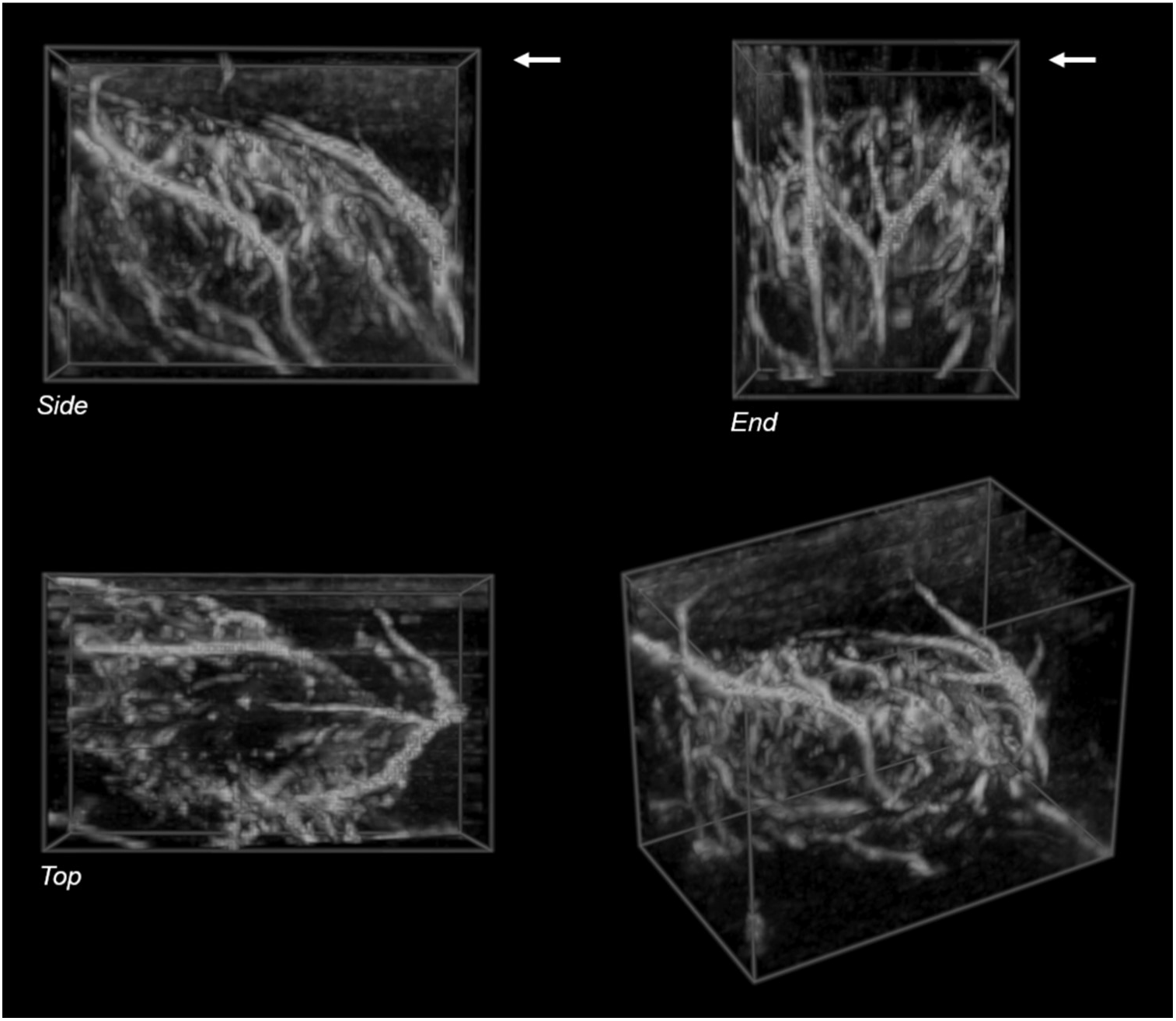
Volume rendering of a 3D reconstruction of Doppler images acquired during a freehand sweep across abscess case ABS1. The color pixels in each 2D frame were segmented and reconstructed without the background gray-scale image. The skin surface is at the upper edge of the Side and End views in the top row (arrows). The bottom left shows an oblique perspective view of the reconstructed volume. Volume size: 33 x 20 x 25 mm (L x W x H). Voxel size: 0.31 mm.

### Serial case study

The series of cross-sectional B-mode images in Figure 3 show the maturation of case ABS2 over a 4-week period. The abscess evolves from a diffuse mass at 1-week post-injection into an organized mass with a well-defined capsule by week 4. Volume renderings of the 3D gray-scale reconstructions (Figure 4) illustrate the morphological development.

**Figure 3.**
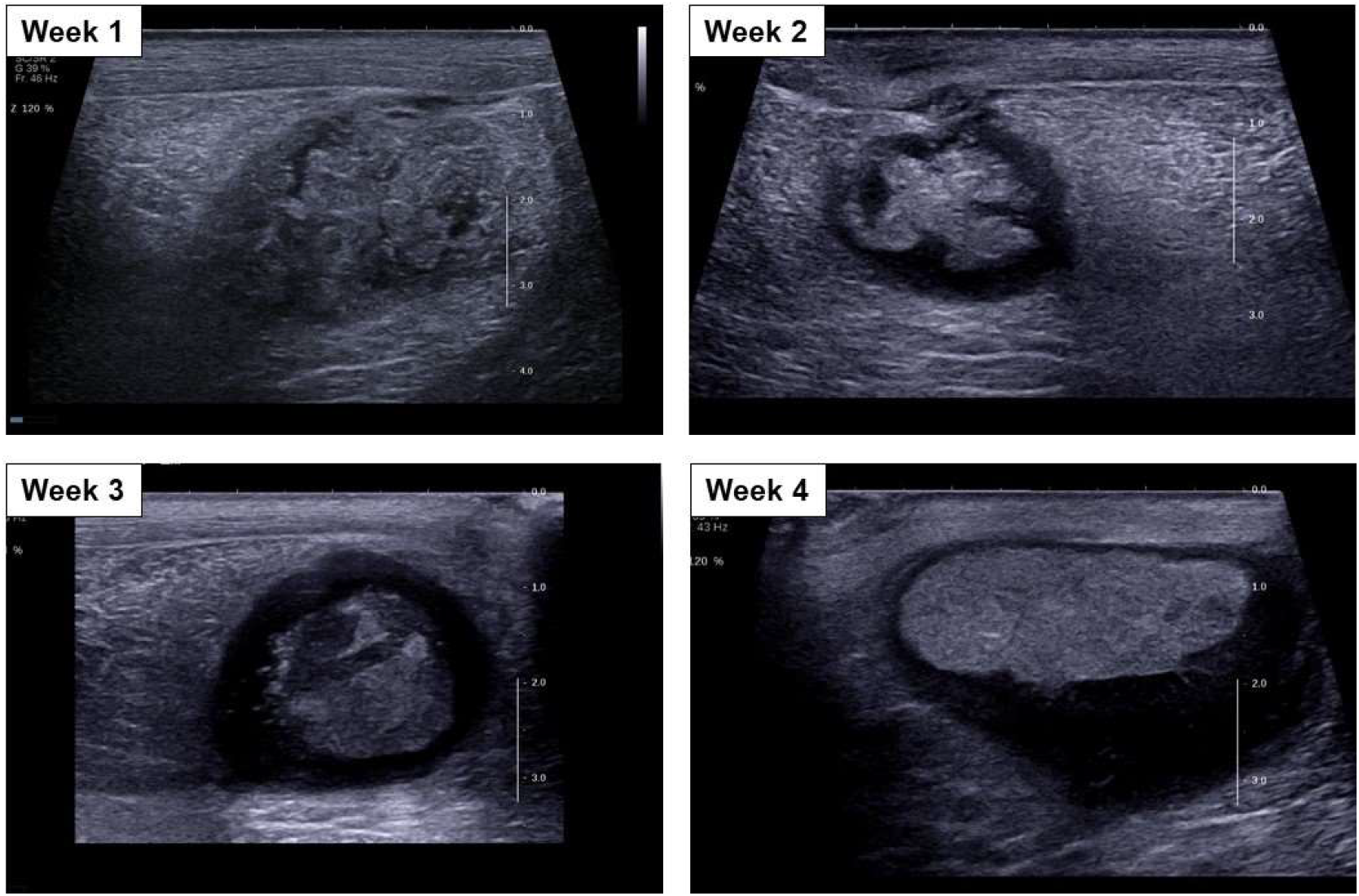
Cross-sectional B-mode images of the intramuscular abscess case ABS2 at weeks 1 through 4 post-injection.

**Figure 4.**
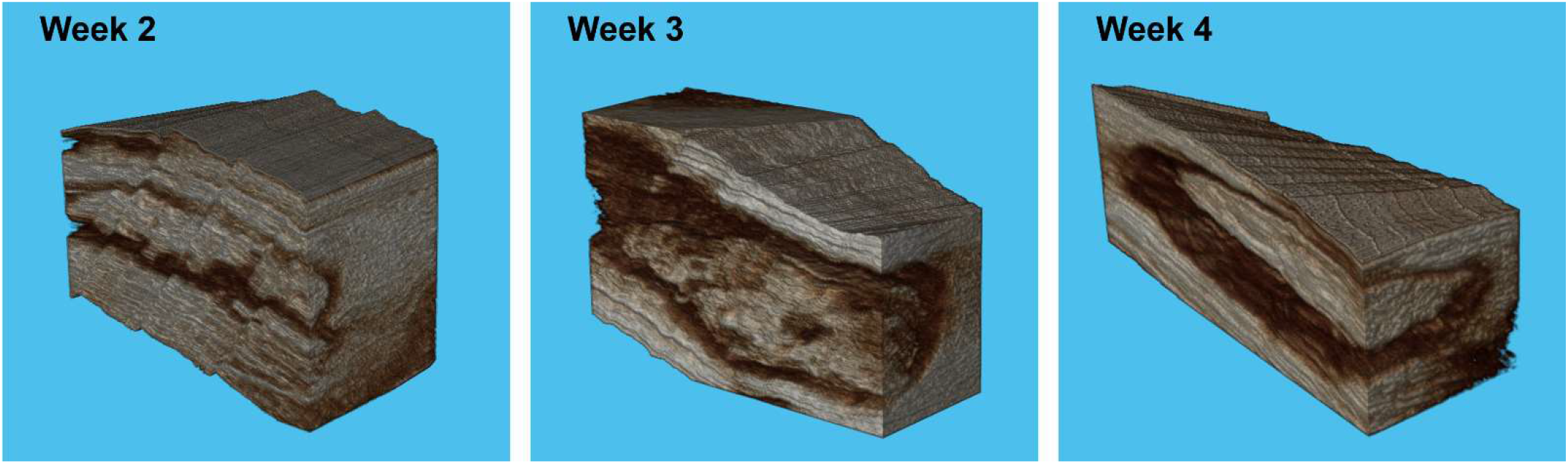
Volume rendering of the 3D volume reconstructions of case ASB2 at weeks 2 through 4 post-injection. The gray-scale volumes are truncated in the cross-sectional and longitudinal directions to visualize the internal structure of the abscess capsule and core. Displayed volume width: 30 mm Week 2, 23 mm Week 3, 32 mm Week 4. Voxel size: 0.08 mm Week 2, 0.16 mm Week 3, 0.32 mm Week 4.

Size and volume changes over time are quantified from the 3D surface reconstructions of the outer extent of the capsule and of the inner core (Figure 5). Table 1 shows the dimensions (Length x Width x Height) and the volume for both the entire abscess and for the abscess core over time. The length of the abscess decreased over time (102 mm at week 2 to 68 mm at week 4) as the volume increased (20.4 ml at week 2 to 37.2 ml at week 4).

**Figure 5.**
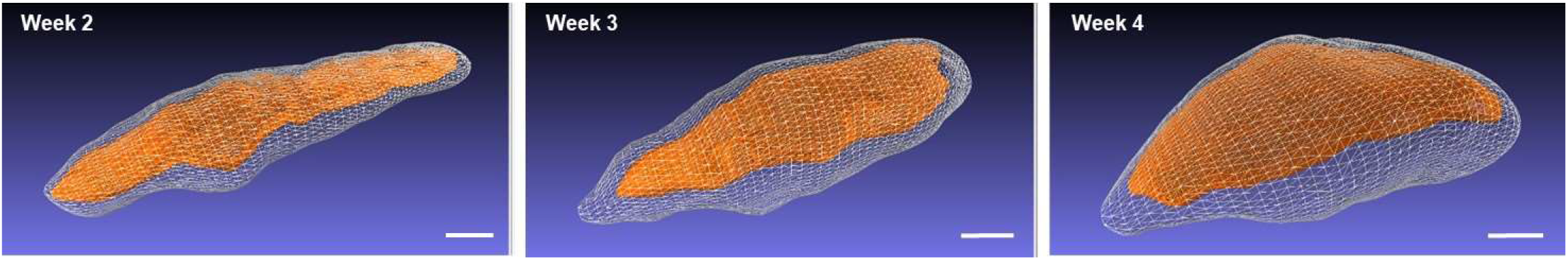
Surface reconstructions generated by manual segmentation of the outer capsule and the inner core for case ABS2 at weeks 2 through 4 post-injection. Scale bars: 1 cm

**Table 1.** 3D Abscess Measurements

| Week | TOTAL |  | CORE |  |  |  |
| --- | --- | --- | --- | --- | --- | --- |
|  | Size (mm) | Volume (ml) | Size (mm) | Volume (ml) | Curvature (1/mm) | Stiffness (kPa) |
| 2 | 102 x 28 x 21 | 20.4 | 99 x 22 x 19 | 8.0 | 0.10 +/- 0.37 | 43.0 ± 19.9 |
| 3 | 88 x 32 x 29 | 30.3 | 80 x 24 x 21 | 12.6 | 0.08 +/- 0.23 | 19.0 ± 11.6 |
| 4 | 68 x 50 x 27 | 37.2 | 63 x 43 x 17 | 16.2 | 0.07 +/- 0.14 | - |

Size: Length x Width x Height Curvature/Stiffness: Mean ± Standard Deviation

Curvature measurements of the core surface over time show a change from an irregular border to a rounded and more well-defined border (Figure 6). Histograms of the curvature measurements at each vertex of the surface mesh (Figure 6) show that the spread in the curvature measurements decreases over time (standard deviation 0.37/mm at week 2 to 0.14/mm at week 4, Table 1) The curvature mean is approximately zero at each time point, indicating a nearly equal distribution of convex and concave curvatures.

**Figure 6.**
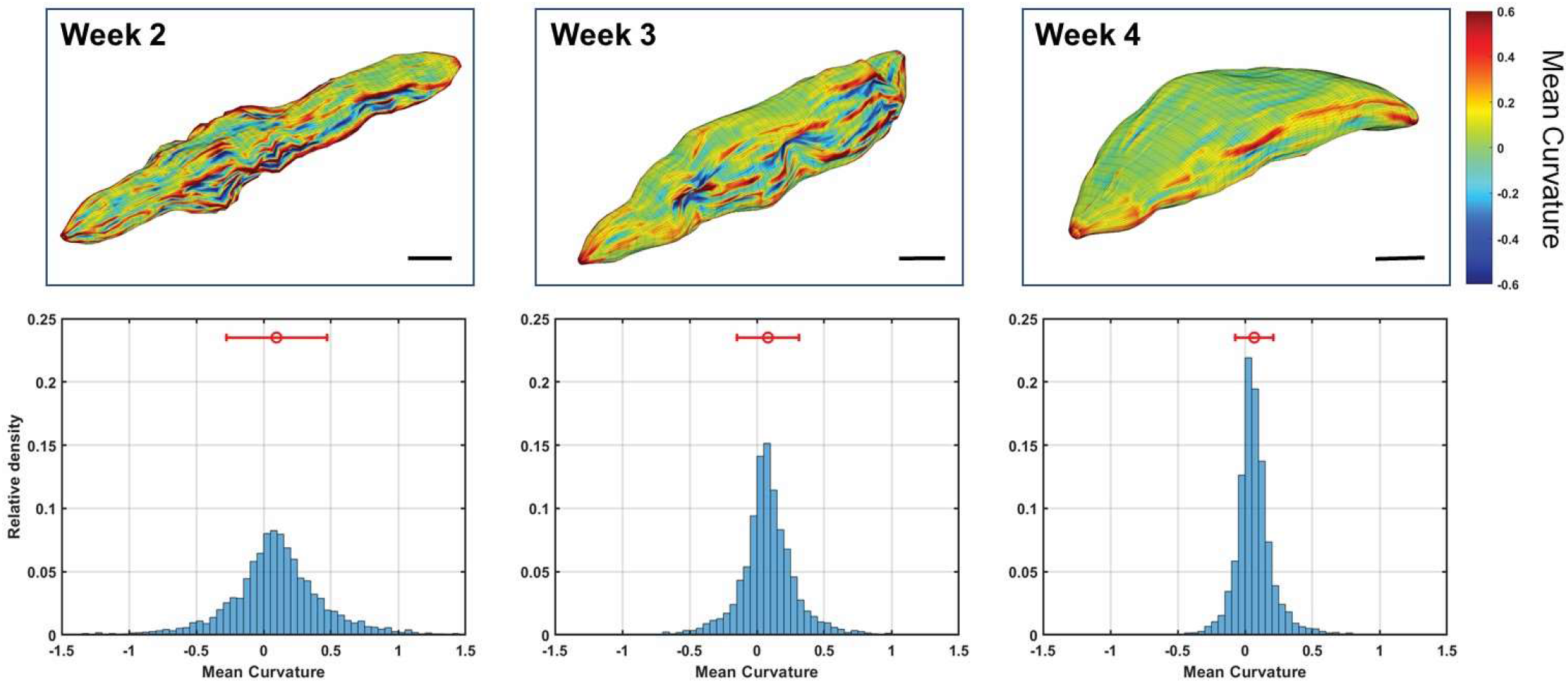
Core surface curvature over time for case ABS2 at weeks 2 through 4 post-injection. Top row: Mean curvature mapped to the surface reconstruction according to the color scale at the right. Scale bars: 1 cm. Bottom row: Histograms of the mean curvature measurements, with the mean and standard deviation shown above. Each histogram is normalized so that the bins sum to one. The curvature measurements of the core surface over time demonstrate a change from an irregular border to a rounded and more well-defined border. Circle: Mean. Bars: Standard deviation. Number of vertices compiled for each histogram: 4558 Week 2, 5144 Week 3, 4622 Week 4.

Cross-sectional SWE images show a reduction in stiffness over the 4-week time period (Figure 7). At Week 4 the abscess core is generally a viscous liquid, with some regions of higher stiffness. 3D transparent volume renderings were produced for the shear-wave measurements in weeks 2 and 3 of the abscess core, based on the untracked cross-sectional sweeps (Figure 8). These 3D renderings show the distribution of stiffness over the entire abscess and illustrate the overall trend toward lower stiffness as the abscess matures and liquefies. This stiffness change is quantified in the histograms in Figure 9, showing the shift toward lower stiffness from week 2 to week 3 (43 kPa to 19 kPa, Table 1).

**Figure 7.**
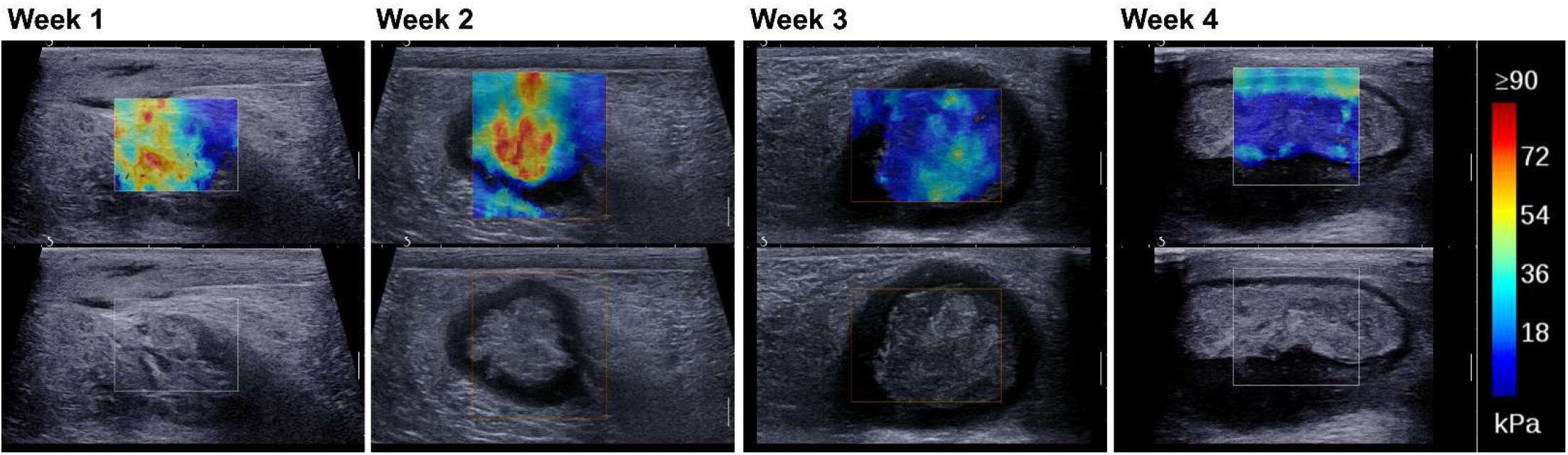
Short-axis Shear Wave Elastography images for case ABS2 at weeks 1 through 4 post-injection showing a decrease in stiffness over time (red to blue).

**Figure 8.**
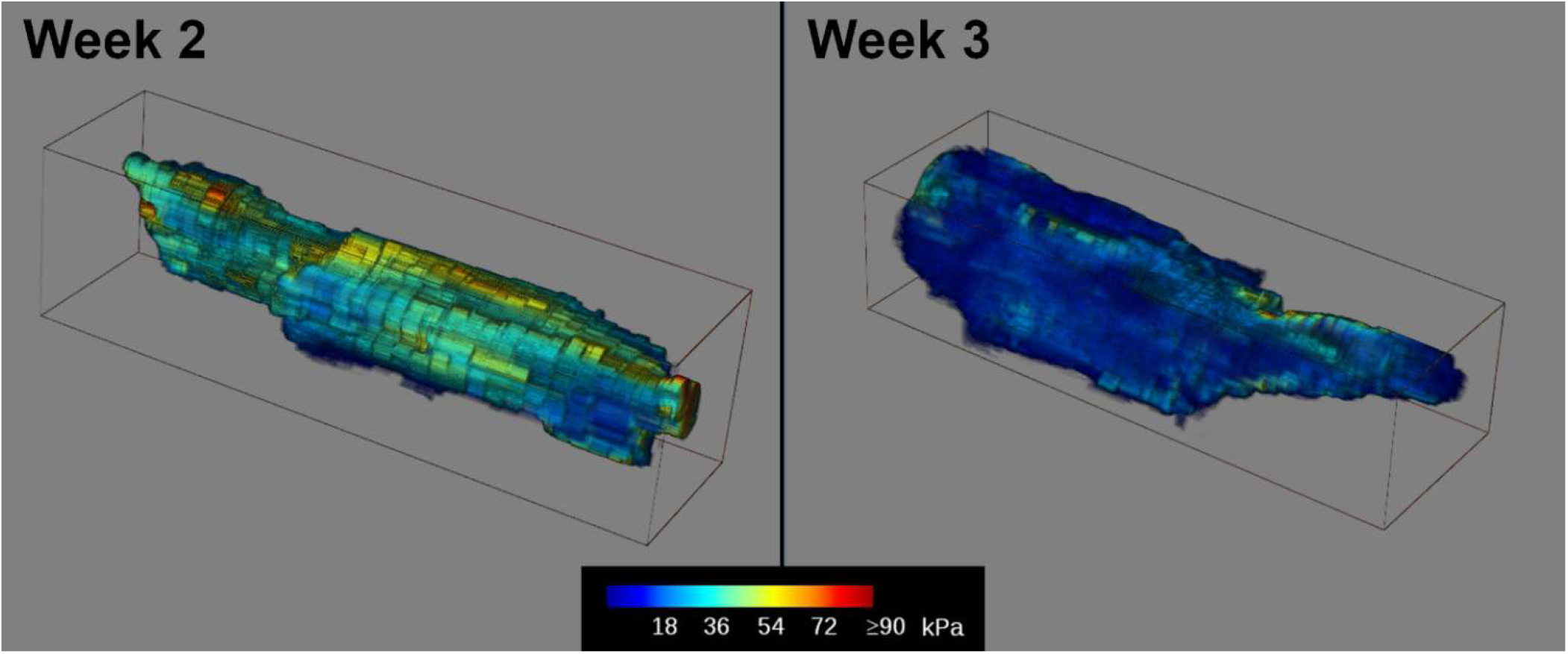
Transparent volume renderings of 3D reconstructions of the SWE measurements in the abscess core at weeks 2 and 3 post-injection for case ABS2. The 3D volumes were generated from untracked freehand scans along the length of the abscess. Voxel colors represent the measured stiffness in kPa according to the color scale shown below. Volume size: 89 x 23 x 25 mm Week 2, 79 x 25 x 19 mm Week 2. Voxel size: 0.08 mm Week 2, 0.07 mm Week 3.

**Figure 9.**
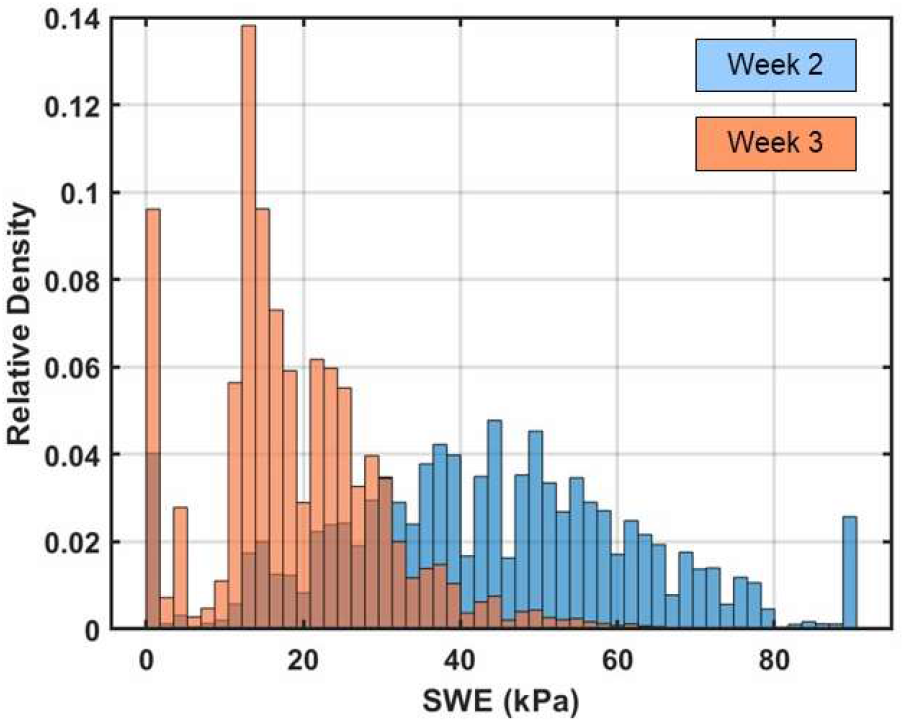
Histograms of elastography measurements compiled over the abscess core at 2 weeks and 3 weeks post-injection for case ABS2. The histograms are normalized so that the bins sum to one. The distribution shifts towards lower stiffness from week 2 to week 3. Number of voxels compiled for each histogram: 28.3 x 10^6^ Week 2, 43.2 x 10^6^ Week 3.

The result of the optical-flow analysis of the compression video is shown in Figure 10a. The yellow arrows show the direction and relative magnitude of the measured displacements; the largest arrow corresponds to 1 mm displacement. Regional differences in stiffness were also observed within the core with SWE (Figure 10b) that correlate with the stationary region around which the internal flow was measured. On histological analysis, this stationary region with higher stiffness was shown to be composed of muscle tissue (Figure 10c).

**Figure 10.**
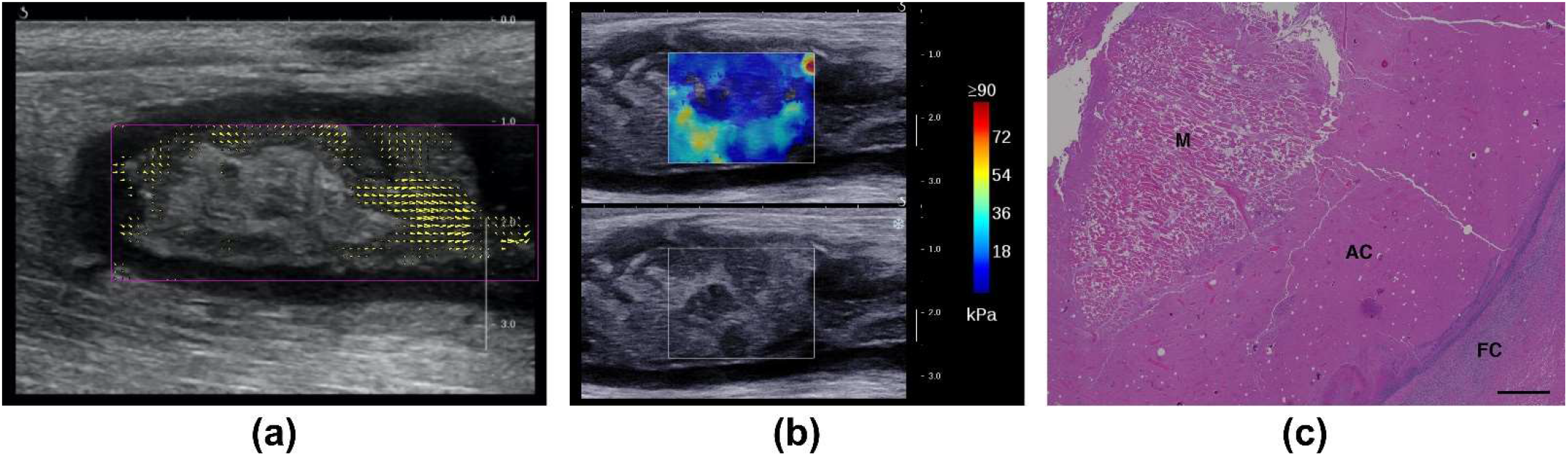
Internal flow with compression and core heterogeneity for the intramuscular abscess case ABS2. (a) Regions of flow with external compression are identified and quantified by comparing regional frame-to-frame displacements at week 3 post-injection. The yellow arrowheads show the direction and magnitude of the displacements. The largest arrows correspond to 1 mm displacements; arrows with magnitudes less than 0.35 mm are not shown. (b) Long-axis SWE image shows regions of varying stiffness in the core at week 3 post-injection. The region with higher stiffness corresponds to the stationary region in (a). (c) Histology of the abscess core with Hematoxylin and Eosin at week 4 post-injection shows a block of muscular tissue (M) within the abscess core (AC). FC: Fibrous Capsule. Scale bar: 1000 microns.

Doppler imaging for case ABS2 showed the development over time of a network of blood vessels within the hypoechoic capsule (Figure 11). At week 4 a dense network of vessels is seen within the capsule, organized around the avascular core. Immunohistochemistry for this case shows the details of the vascular network on both the deep and lateral sides of the capsule that matched the Doppler images (Figure 12). No blood vessels were observed within the abscess core.

**Figure 11.**
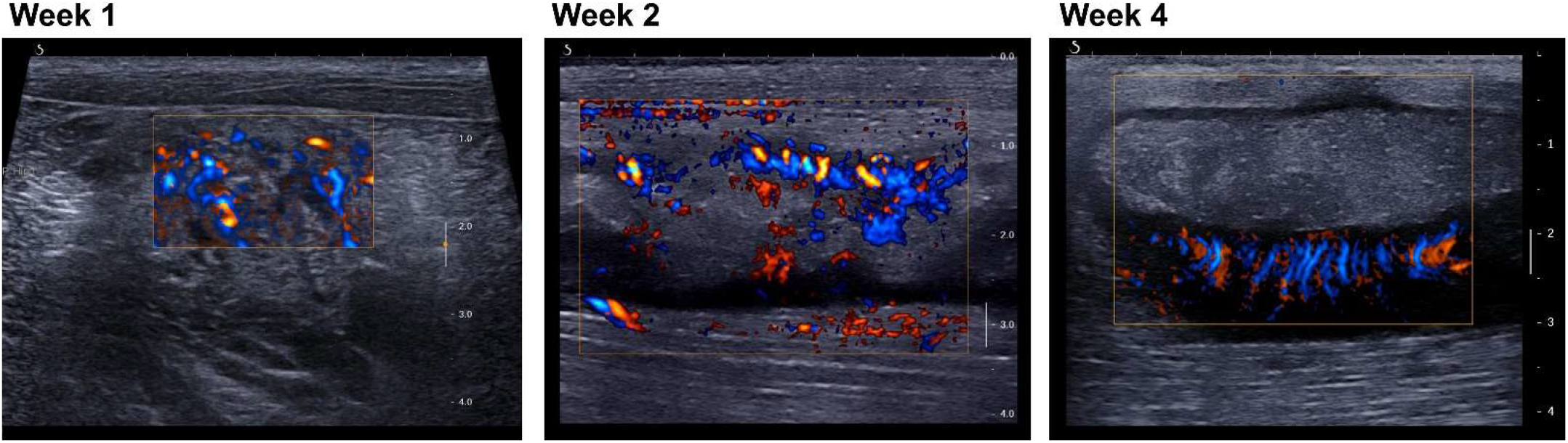
Long-axis views of abscess case ABS2 with color Doppler imaging show a network of vessels developing from a diffuse pattern throughout the injection site (week 1) to an organized network within the capsule surrounding the avascular core (week 4). Week 1: standard directional color power Doppler. Weeks 2 and 3: plane-wave directional color power Doppler.

**Figure 12.**
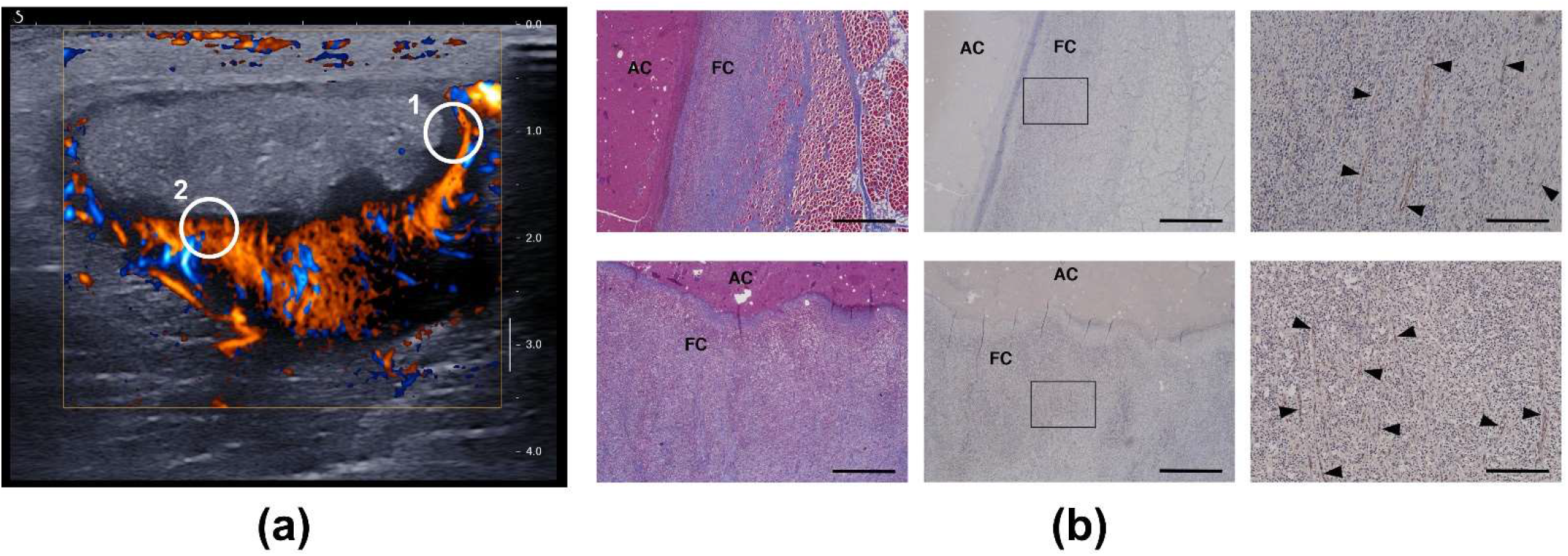
(a) Short-axis plane-wave directional color Doppler image (4 weeks post-injection) of the vascularized capsule and avascular core of case ABS2. The circled regions indicate the general locations for which histology images were obtained. (b) Histology of the capsule confirms a network of vessels with circumferential orientation on the lateral side (top row, circle 1) and oriented toward the core on the deep side (bottom row, circle 2). The vessels are indicated by the black arrow heads in the high-magnification images in the right column. Left: Masson’s trichrome. Middle and right: CD31 labelled. Right: magnification of the boxed regions in the middle column. FC: Fibrous Capsule. AC: Abscess Core. Scale bar: 1000 microns (left and middle) / 200 microns (right).

## Discussion

As an abscess matures from a region of inflammation (phlegmon) to an encapsulated organized collection of viscous pus, it consolidates and a fibrous capsule is formed around the purulent material. The maturation process includes angiogenesis of permeable vessels in the capsule, through which inflammatory cells (neutrophils, lymphocytes, macrophages) enter the infected area. Subsequently the abscess core liquefies as the inflammatory cells clear the infection, and the abscess consolidates and retracts.

In our animal model, the injection of bacteria and dextran initially resulted in the development of a phlegmon that then evolved into an organized mature abscess over 2-4 weeks. B-mode ultrasound allowed us to identify key structural characteristics, namely a well-defined hypoechoic capsule surrounding a core with mixed echogenic material, which were used as an indicator of maturity (Lu et al. 2019). Doppler imaging was used to further evaluate abscess maturity, with mature abscesses having rich vasculature within the hypoechoic capsule, and an avascular core.

In general, the pig abscesses were more echogenic than often observed in humans, although echogenicity of human abscesses can vary (Kuligowska et al. 1982). One contributing factor may be the addition of dextran particles (~150 μm) to the inoculate, which likely increases scattering and makes the abscess appear brighter. The purpose of the dextran was to elicit a persistent inflammatory response to aid in the reliable generation of an abscess (Wang et al. 2020).

The surface reconstructions from 3D B-mode scans allow quantification of size, volume, and core curvature over time. Surface curvature analysis of the core shows an organizational change in configuration as the abscess matures. The reduced spread of the curvature measurements over time signify a reduction sharp bends in the surface and a general trend toward a more smooth and rounded shape. Similar curvature analysis has been used to describe changes in surface complexity in the developing fetal brain (Hu et al. 2013), and to distinguish between different kidney stone types (Duan et al. 2013). These abscess curvature changes match the development over time previously described for liver abscesses (Kunze et al. 2015).

Shear wave elastography shows reduction in core stiffness during abscess maturation (Figure 7). The 3D volume reconstructions of SWE scans at weeks 2 and 3 quantify this stiffness change, and demonstrate that the change is generally observed throughout the abscess core (Figure 8).

The internal flow observed at week 3 indicates the presence of lower-viscosity liquid components in the core. Internal flow with compression has been noted as a component of the clinical ultrasound examination of abscesses that helps distinguish an abscess from other conditions, such as cellulitis and phlegmon (Adhikari and Blaivas 2012). SWE at this site also demonstrated the heterogeneity of the abscess core. Histology confirmed that the region with higher stiffness and lack of flow with compression was residual muscle tissue, distinct from the general viscous content of the abscess core.

Doppler imaging shows development of a vascular capsule with an avascular core, which were confirmed by histology. This vascularized capsule should correspond to the enhancing rim cited as one of the CT criteria to suggest an abscess (Mavilia et al. 2016). Abscess rim enhancement has also been observed with contrast-enhanced ultrasound (Liu et al. 2008, Kunze et al. 2015).

## Conclusion

Mature abscesses were characterized by an organized, avascular center of varying echogenicity surrounded by a hypoechoic capsule that was highly vascularized on Doppler imaging. The distinction of abscesses from other pathologies is required for planning the proper approach to treatment. In addition, identification of abscess maturity is of clinical relevance because the core must be relatively liquefied in order for drainage to be successful. The measurements of abscess morphology, stiffness and vascularity demonstrated in this study represent ultrasound imaging features that can potentially identify lesions and their suitability for treatment.

## Acknowledgements

Work supported in part by NIH grants 5R01EB019365, R01GM122859 and K01 DK104854.

## Notes

### Competing Interest Statement

The authors have declared no competing interest.

